# Knowledge and Attitude Towards Palliative Care Among Clinical Undergraduate Students of Physiotherapy in Nigerian Universities: A Cross-Sectional Study

**DOI:** 10.1101/2022.06.12.495785

**Authors:** Marufat Oluyemisi Odetunde, Jessie Chiamaka Nwankwo, Afolabi Muyiwa Owojuyigbe, Olurotimi I. Aaron, Adaobi Magraet Okonji

## Abstract

**Introduction:** Palliative care demands a special skill from health care providers which begins with the right attitude and adequate knowledge during training. Little is known about knowledge and attitude of physiotherapy undergraduates towards palliative care. Objective of this study was to assess knowledge and attitude of clinical undergraduates of physiotherapy in Nigerian universities towards palliative care.

**Method:** This was a cross sectional survey involving 281 (132 females and 149 males) clinical physiotherapy undergraduates from 10 purposively selected universities across Nigeria. Palliative care knowledge was assessed with a structured questionnaire while attitude was assessed using FROMMELT attitude towards care of the dying scale. Both questionnaires were administered online via Google forms. Socio-demographic data of each respondent was also collected. Data were analyzed using descriptive and inferential statistics. Level of significance was set at p<0.05.

**Results:** Majority of respondents were males (53%) aged 21-25 years (87.9%) from Obafemi Awolowo University (40.3%) and had above average knowledge (62%). Almost all, (99.6%) of the respondents displayed favourable attitude. There were significant associations between knowledge and university of respondents (p=0.001), attitude and age of respondents (p=0.003), as well as attitude and geopolitical zone of origin of respondents (p=0.006). There was significant correlation between knowledge and attitude of respondents (r =0.441, p =0.001).

**Conclusion:** There are knowledge gaps, but positive attitudes towards palliative care among clinical physiotherapy undergraduates in Nigeria. This signifies an essential and urgent need for inclusion, reform or improvement in palliative care education in the curriculum of undergraduate physiotherapy training in Nigerian universities.

## Introduction

The number of patients with cancer and other terminal diseases continue to rise in the last decades leading to rapid growth in concern for severely ill and dying patients [1]. Palliative Care (PC) is a method that increases the quality of life of patients with life-threatening illness and their families through prevention and relief of distress by early identification, comprehensive assessment and treatment of pain and other physical, psychosocial and spiritual problems [2]. PC comprises an interdisciplinary team of experts including physicians (specialists in hospice and palliative medicine), nurses, specialty advanced practice nurses and registered nurses, social workers, chaplains, pharmacists, rehabilitation therapists, direct care workers, and family members [3]. PC is required for a wide range of chronic diseases which defy curative treatment. These include cardiovascular diseases, cancer, chronic respiratory diseases, AIDS, diabetes, kidney failure, chronic liver disease, multiple sclerosis, rheumatoid arthritis, neurological diseases, dementia, congenital anomalies and drug resistant tuberculosis [4].

Patients with terminal illnesses often experience significant symptom burden which may result from various complications while treatments can also cause significant side effects, all of which significantly decrease quality of life (QOL) of patients [3]. In both economically advanced and economically less developed countries, people live and die with uncontrolled pain and physical symptoms, unresolved psychosocial and spiritual problems, fears and loneliness and complicated bereavement, which could be ameliorated with PC [5,6].

Palliative care is established and mostly provided in high-income countries such as the United States of America in over 60% of hospitals. Whereas 80% of the global need of PC lies in in low- and middle-income countries [7], yet it is just an emerging medical specialty established in Asia and few countries in Africa [8]. PC is still relatively uncommon in Nigeria as there are only a few centres across the country where PC services are being rendered. Ineffectively implemented policies of the Hospice and PC Association of Nigeria (HPCAN), and administrative bureaucracy of the Nigerian government are the challenges hindering implementation of effective PC services in Nigeria [9]. These polices seem non-existent owing to many challenges including lack of standard facilities and equipment, lack of professionals to non-inclusion of some professionals, low awareness and knowledge, difficulty communicating about cancer diagnosis and management with patients, limited availability of medications including oral opioids [9]. All these have diverse effects on the knowledge of health care professionals in training as they have limited hands on experience with cases that require PC [9] and deficient in training at the undergraduate level. In Nigeria, only two Universities offer PC; the University of Ibadan as revised undergraduate curriculum; while University of Ilorin offers it in a post-graduate diploma. Many just operate workshops and continuous professional training [6].

Palliative Care education in economically advanced world indicated that 28 European countries have palliative medicine in the curriculum of at least one medical university, while it is taught in all medical schools in 13 countries; whereas in 14 countries palliative medicine was included in the under-graduate medical curriculum [10]. Higher level education in palliative care is now available in several countries in Africa, including South Africa, Uganda, and Tanzania [11]. The Institute of Hospice and Palliative Care in Africa in partnership with Makerere University, also provides training to healthcare professionals across Africa [11]

Professionals equipped with basic PC skills can deliver effective front-line supportive care [3]. The role of each healthcare professional in the delivery of PC to terminally ill patients is uniquely distinct and highly important, while an effective PC service delivery requires an informed health sector [12]. The roles become even more important with increasing need for PC as a result of the ageing of populations and the rising burden of cancer, non-communicable diseases and some communicable diseases. Professionals in PC team should have appropriate level of education and competence in handling the varying needs and requirements of PC. This necessitates evidence-based PC knowledge and education level as well as training needs of these professionals. [13]

Although available evidence supports the role of physiotherapists and rehabilitation in PC, literature describing knowledge, role and view of physiotherapy about palliative care are scarce [14, 15, 16]. Physiotherapy in PC aims to maintain QOL by relieving stress from pain and side effects of treatment delivered at four levels of prevention, pre and post-operative care, acute institutional and community-based rehabilitation [16]. Thus, appropriate referral to PC physiotherapy is critical for optimal and patient-centred care. Education and training in PC impacts the level of care provided and team participation of physiotherapists. Thus PC education is important for physiotherapy students at undergraduate level to train competent and responsible members of PC team in the society who lead physiotherapy profession successfully by providing useful information and realistic implementation [17]. Little is known about knowledge of PC among physiotherapists in Nigeria at all levels of intervention. Therefore, specific objective of this study was to assess knowledge and attitude of clinical undergraduate students of physiotherapy in Nigerian Universities towards PC. We hypothesized that there would be no significant correlation between knowledge and attitude Nigerian physiotherapy undergraduates towards PC.

## Methods

This cross sectional study was a web-based survey involving clinical undergraduate physiotherapy students in Nigerian universities where physiotherapy is offered as a course at undergraduate level. The target population for this study was a total of 1,152 clinical physiotherapy students across the universities. The number of students in each school at the time of data collection were as follows, 214 students in Obafemi Awolowo University, Ile-Ife, Osun state, 90 students in University of Ibadan, Oyo state, 80 students in University of Lagos, Lagos state, 95 students in Bowen University, Iwo, Osun state, and 67 students in University of Medical Sciences Ondo, Ondo state (all in southwestern Nigeria); 145 students in University of Nigeria Nsukka, 130 students in Nnamdi Azikiwe University Anambra state (southeastern Nigeria), 72 students in University of Benin (south-southern Nigeria); 157 students in Bayero University, Kano, Kano state, 102 students in University of Maiduguri, Borno state (northern Nigeria). Respondents were included if they were in clinical years of training and in clinical postings within or outside their respective university teaching hospitals. Exclusion criteria were students in pre-clinical years, clinical students on leave of absence or those not in clinical postings for other reasons.

Using sample size table based on desired accuracy with confidence level of 95% [18], Population size from table-1000 (range of 1000-1499), margin error E - 5% and, variance of the population set at P - 50%, sample size n was given as 278. Hence 278 respondents were required for this study. A total of 281 respondents (132 females and 149 males) completed the questionnaire giving a complete (100%) response rate.

The instrument for data collection was a structured questionnaire [19] (Appendix I) and the FROMMELT attitude towards care of the dying (FATCOD) scale [12] (Appendix II). The questionnaires were presented in three sections (A, B and C). SECTION A contained information on socio-demographic details of respondents, SECTION B was a 39-point questionnaire with 10 sub-divisions on knowledge of PC, and SECTION C was a 30-item assessment of attitude towards PC, the FATCOD scale. Respondents were to choose ‘yes’ or ‘no’ for answers in section B, where total score was obtained by adding each correct response from each item on the questionnaire [19]. In section C, responses were rated on a 5-point Likert scale; ranging from strongly disagree to strongly agree. Scoring was done using basic additions, percentages of right and wrong answers and fractions as utilized in previous studies [12]. Detailed explanation and instruction about this study was given to respondents and their written informed consent was obtained. Anonymity was strictly observed as name of respondents were not requested. Respondents were assured there were no right or wrong answers, non-maleficent of the study and freedom to refuse to respond to the questionnaire or withdraw from the study at any stage without any penalty. The research protocol was approved by appropriate Health and Research Ethics Committee. Respondents from Obafemi Awolowo University, Ile-Ife, who met the inclusion criteria, self-completed the questionnaire by paper and pen. For respondents in the other nine universities, questionnaire link of Google forms was shared with the representative of each class who in turn sent it to the class WhatsApp groups. The National Association Physiotherapy students in Nigeria had their annual convention in 2021. Required information about clinical physiotherapy students in each University was obtained from the representative of each University at the convention. Ample time was given to complete the questionnaire, which was submitted online after completion over a period of two months from August to September 2021.

## Data analysis

Descriptive statistics of frequencies and percentages were used to summarize the socio-demographic variables of respondents. Chi square test was used to assess the association between each of knowledge and attitude with socio-demographic variables. Pearson’s correlation method was used to test correlation between knowledge and attitude. Data was analyzed using Statistical Package for Social Sciences (SPSS) software version 16. The level of significance was set at p< 0.05.

## Results

Majority of the respondents were males (53%) at 400 level (4^th^ year) of study (50.9%), aged 21-25 (87.9%) with only 6.8% aged 20 years and below, Christians (73.7%) and from south-western Nigeria (61.9%). Frequency distribution of respondents by institution is shown in figure 1. Distribution of respondents’ knowledge about PC is presented in Figure 2. Over half (62%) of respondents had above average knowledge whereas almost all (99.6%) of respondents displayed favourable attitude towards PC.

**Figure 1:**
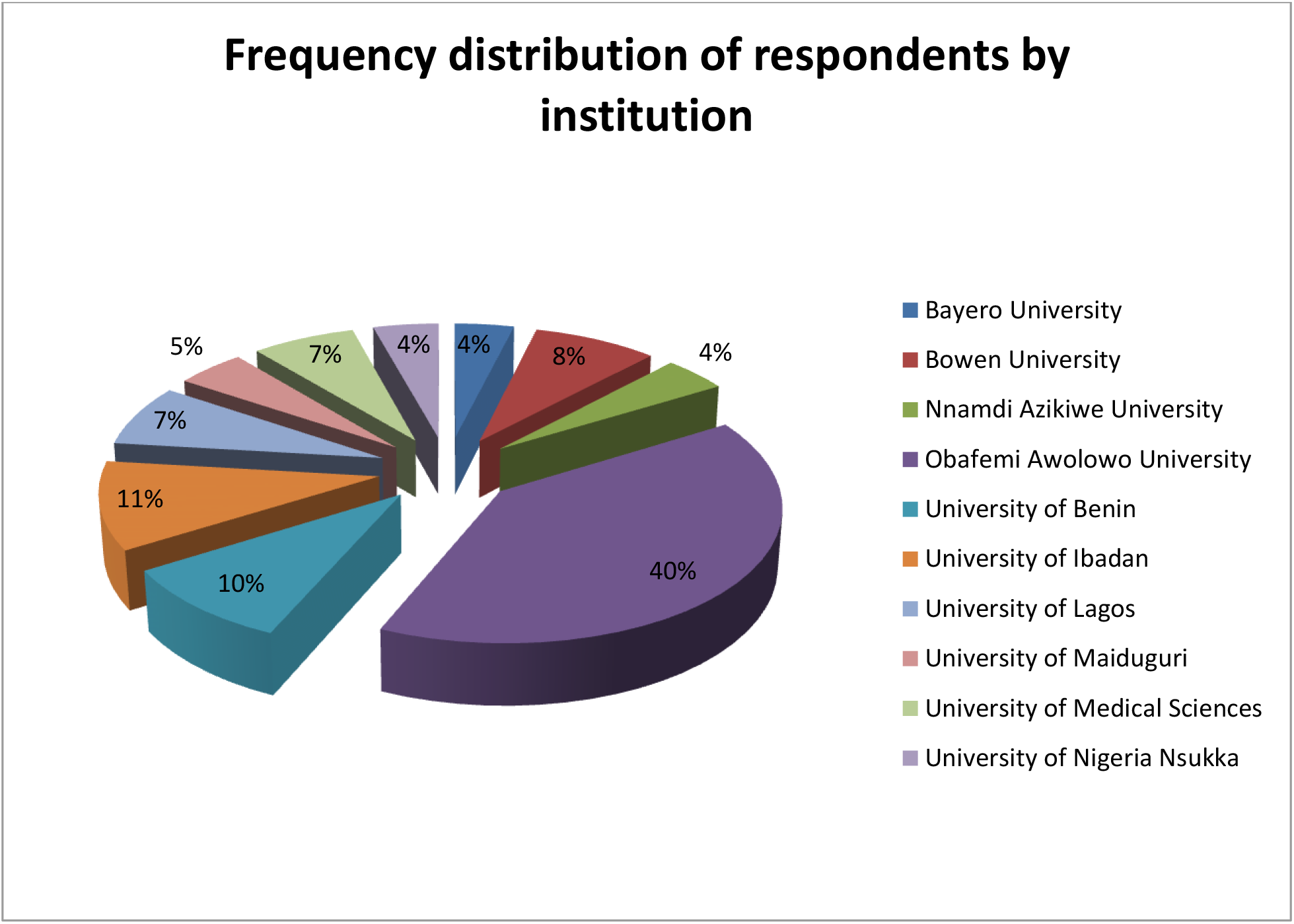
Distribution of clinical undergraduate students’ of physiotherapy in Nigeria by University.

**Figure 2:**
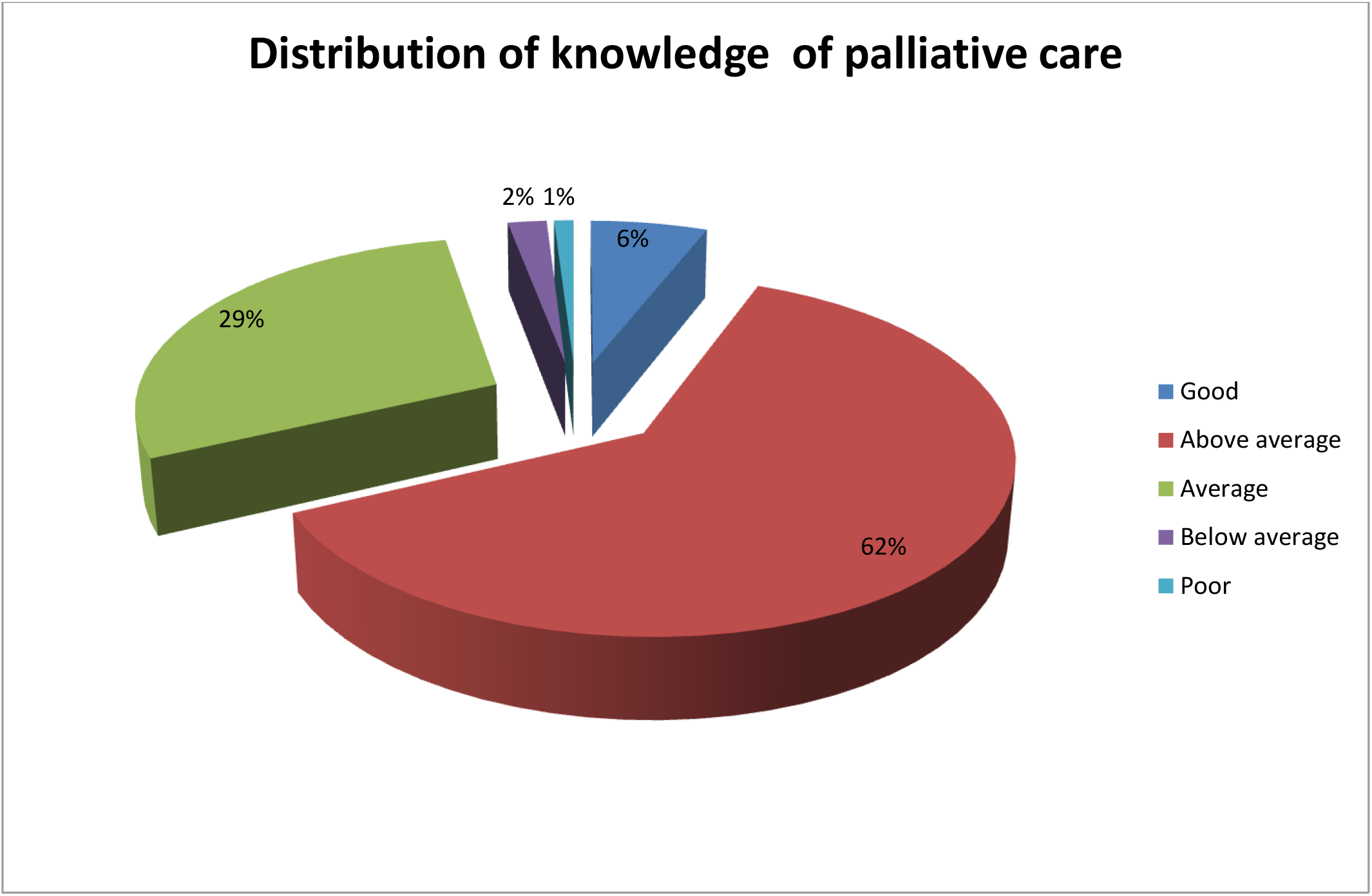
Distribution of knowledge about PC among clinical undergraduate students’ of physiotherapy in Nigeria.

Respondents’ knowledge about PC revealed that 24 out of a total of 39 items on knowledge are with expected positive (―yes‖) answers. Over 50% of respondents chose the correct responses for 29 out of 39 items, whereas less than 50% of the respondents chose incorrect responses for 28 out of 39 items. Majority (72.6%) of respondents correctly chose ‘yes’ that PC is active care of the dying while 85.4% wrongly chose ‘yes’ that PC hastens death, whereas 85.1% chose ‘’yes’ that PC is needed for metastatic cancer with uncontrolled pain. 87.9% correctly chose ‘’no’ that morphine causes death in all dying patients, 78.3 correctly chose ‘’yes’ that common side effects of morphine in PC setting is drowsiness, common non-pain symptoms encountered in PC was correctly chosen as ‘yes’ to be delirium by 72.2% of respondents, whereas respondents generally correctly chose ‘yes’ for all components of dying while PC multidisciplinary team was wrongly chosen as ‘no’ to comprise radiotherapist by the least (32.7%) number of respondents. Respondents’ most preferred choices on attitude towards PC indicated that over half of them agreed with the most preferred choices on seven out of the 30 attitude items viz ‘there are times when death is welcomed by the dying person’ (56.2%), ‘family should be involved in the physical care (feeding, personal hygiene) of the dying person’ (52.3%), ‘families need emotional support to accept the behaviour changes of the dying person’ (50.2%), ‘families should maintain as normal an environment as possible for their dying member’ (51.2%), ‘it is beneficial for the dying person to verbalize his or her feelings’(52.0%), ‘care should extend to the family of the dying person’(55.2%), and that ‘caregivers should permit dying persons to have flexible visiting schedules’(54.1%). All other attitude items were rated by less than 50% of respondents as most preferred choices.

There was moderate and significant correlation between knowledge and attitude of respondents towards PC (r =0.441, p-value =0.001). Table 1 shows the association between knowledge and socio-demographic variables of respondents. The results indicated that there was a significant correlation between knowledge of PC and University of respondents (p-value=0.001). Respondents’ association between attitude and socio-demographic variables are shown in table 2. There were significant correlation between attitude towards PC and age of respondents (p-value =0.003) and also between attitude towards PC and tribes (geopolitical zone of origin) of respondents (p-value=0.006)

**Table 1:** Association between Knowledge and Socio-demographic Variables of Respondents.

| Variables | Good | Above average | Average | Below average | Poor | Total n (N) | $\chi^2$ | p-value |
| --- | --- | --- | --- | --- | --- | --- | --- | --- |
| **Gender** |  |  |  |  |  |  |  |  |
| Female | 9(6.82) | 88(66.7) | 29(22.0) | 2(1.52) | 4(3.03) | 132 | 9.995 | 0.41 |
| Male | 7(4.70) | 87(58.4) | 52(35.0) | 3(2.01) | 0(0.00) | 149 |  |  |
| **Level of study** |  |  |  |  |  |  |  |  |
| 400 | 6(4.20) | 84(58.7) | 51(35.6) | 0(0.00) | 2(1.40) | 143 | 11.639 | 0.200 |
| 500 | 10(7.25) | 91(66.0) | 30(21.7) | 5(3.62) | 2(1.45) | 138 |  |  |
| **Age (years)** |  |  |  |  |  |  |  |  |
| 20 and below | 1(5.26) | 8(42.1) | 9(46.4) | 0(0.00) | 1(5.26) | 19 | 15.810 | 0.200 |
| 21-25 | 15(6.07) | 161(65.2) | 64(26.0) | 4(1.62) | 3(1.21) | 247 |  |  |
| 26-30 | 0(0.00) | 6(42.9) | 7(50.0) | 1(7.14) | 0(0.00) | 14 |  |  |
| Above 30 | 0(0.00) | 0(0.00) | 1(100) | 0(0.00) | 0(0.00) | 1 |  |  |
| **Religion** |  |  |  |  |  |  |  |  |
| Christianity | 12(5.80) | 129(62.3) | 60(29.0) | 3(1.45) | 3(1.45) | 207 | 3.293 | 0.915 |
| Islam | 4(6.35) | 40(63.49) | 16(25.4) | 2(3.17) | 1(1.59) | 63 |  |  |
| Other | 0(0.00) | 6(54.5) | 5(45.5) | 0(0.00) | 0(0.00) | 11 |  |  |
| **Geopolitical zone of tribe** |  |  |  |  |  |  |  |  |
| North central | 0(0.00) | 7(100) | 0(0.00) | 0(0.00) | 0(0.00) | 7 | 93.151 | 0.149 |
| North-east | 0(0.00) | 2(100) | 0(0.00) | 0(0.00) | 0(0.00) | 2 |  |  |
| North-west | 1(4.16) | 10(4.16) | 12(50.0) | 1(4.16) | 0(0.00) | 24 |  |  |
| South-south | 0(0.00) | 9(50.0) | 7(38.9) | 0(0.00) | 2(11.1) | 18 |  |  |
| South-east | 4(7.27) | 32(58.2) | 15(27.3) | 2(3.64) | 2(3.64) | 55 |  |  |
| South-west | 11(6.32) | 114(65.5) | 47(27.0) | 2(1.15) | 0(0.00) | 174 |  |  |
| Foreign | 0(0.00) | 1(100) | 0(0.00) | 0(0.00) | 0(0.00) | 1 |  |  |
| **Institution** |  |  |  |  |  |  |  |  |
| Bayero University | 0(0.00) | 7(63.6) | 4(36.4) | 0(0.00) | 0(0.00) | 11 (157) | 74.781 | 0.001 |
| Bowen University | 2(8.70) | 15(65.2) | 4(17.4) | 2(8.70) | 0(0.00) | 23 (95) |  |  |
| Nnamdi Azikiwe University | 0(0.00) | 10(83.33) | 1(8.33) | 0(0.00) | 1(8.33) | 12 (130) |  |  |
| Obafemi Awolowo University | 9(7.96) | 82(72.6) | 22(19.5) | 0(0.00) | 0(0.00) | 113 (214) |  |  |
| University of Benin | 1(3.70) | 12(44.4) | 13(48.1) | 0(0.00) | 1(3.70) | 27 (72) |  |  |
| University of Ibadan | 1(3.33) | 21(70.0) | 8(26.7) | 0(0.00) | 0(0.00) | 30 (90) |  |  |
| University of Lagos | 1(4.76) | 10(47.6) | 9(42.9) | 1(4.76) | 0(0.00) | 21 (80) |  |  |
| University of Maiduguri | 0(0.00) | 5(38.5) | 7(63.8) | 1(7.70) | 0(0.00) | 13 (102) |  |  |
| University of Medical Sciences | 2(10.5) | 8(42.1) | 9(47.4) | 0(0.00) | 0(0.00) | 19 (67) |  |  |
| University of Nigeria Nsukka | 0(0.00) | 5(41.7) | 4(33.3) | 1(8.33) | 2(16.7) | 12 (145) |  |  |

**Table 2:** Association between Attitude and Socio-demographic variables of Respondents.

| **Variables** | **Favourable**<br>**Frequency (%)** | **Unfavourable**<br>**Frequency (%)** | **Total**<br>**n (N)** | **$\chi^2$** | **p-**<br>**value** |
| --- | --- | --- | --- | --- | --- |
| **Gender** |  |  |  |  |  |
| Female | 131(99.2) | 1(0.76) | 132 | 1.133 | 0.287 |
| Male | 149(100) | 0(0.00) | 149 |  |  |
| **Level of study** |  |  |  |  |  |
| 400 | 142(99.3) | 1(0.70) | 143 | 0.968 | 0.325 |
| 500 | 138(100) | 0(0.00) | 138 |  |  |
| **Age (years)** |  |  |  |  |  |
| 20 and below | 18(94.7) | 1(5.26) | 19 | 13.839 | 0.003 |
| 21-25 | 247(100) | 0(0.00) | 247 |  |  |
| 26-30 | 14(100) | 0(0.00) | 14 |  |  |
| Above 30 | 1(100) | 0(0.00) | 1 |  |  |
| **Religion** |  |  |  |  |  |
| Christianity | 207(100) | 0(0.00) | 207 | 3.473 | 0.176 |
| Islam | 62(98.4) | 1(1.60) | 63 |  |  |
| Other | 11(100) | 0(0.00) | 11 |  |  |
| **Geopolitical zone of tribe** |  |  |  |  |  |
| North central | 7(100) | 0(0.00) | 7 | 39.283 | 0.006 |
| North-east | 2(100) | 0(0.00) | 2 |  |  |
| North-west | 24(100) | 0(0.00) | 24 |  |  |
| South-south | 17(94.4) | 1(5.56) | 18 |  |  |
| South-east | 55(100) | 0(0.00) | 55 |  |  |
| South-west | 174(100) | 0(0.00) | 174 |  |  |
| Foreign | 1(100) | 0(0.00) | 1 |  |  |
| **Institution** |  |  |  |  |  |
| Bayero University | 11(100) | 0(0.00) | 11 (157) | 9.441 | 0.398 |
| Bowen University | 23(100) | 0(0.00) | 23 (95) |  |  |
| Nnamdi Azikiwe University | 12(100) | 0(0.00) | 12 (130) |  |  |
| Obafemi Awolowo<br>University | 113(100) | 0(0.00) | 113 (214) |  |  |
| University of Benin | 26(96.3) | 1(3.70) | 27 (72) |  |  |
| University of Ibadan | 30(100) | 0(0.00) | 30 (90) |  |  |
| University of Lagos | 21(100) | 0(0.00) | 21 (80) |  |  |
| University of Maiduguri | 13(100) | 0(0.00) | 13 (102) |  |  |
| University of Medical<br>Sciences | 19(100) | 0(0.00) | 19 (67) |  |  |
| University of Nigeria<br>Nsukka | 12(100) | 0(0.00) | 12 (145) |  |  |

## Discussion

This study assessed knowledge and attitude towards PC among clinical undergraduates of physiotherapy in Nigerian universities. Two hundred and eighty-one students responded to the online survey. Above average knowledge of PC was demonstrated by majority (62%) of the respondents while only 6% had good knowledge. This appears better than inadequate basic PC knowledge among healthcare undergraduates in India [12] and deficiencies in understanding various concepts of palliative care among students reported by Sujatha et al., [20]. These gaps in the knowledge of PC among the current undergraduate sample may be as a result of deficiency in the PC education curriculum and training received by these students at undergraduate level. Findings from the previous studies in other countries demonstrated widespread deficiencies in understanding PC and its philosophy, pain and non-pain symptom assessment, understanding resuscitation in PC setting, communication, and interdisciplinary care for patients and their families [12]. In light of the above, although better than other climes, this study also depicts some deficiencies in basic aspects of PC knowledge among undergraduate clinical physiotherapy students in Nigeria.

About three-quarter (72.6%) respondents were able to relate active care of dying with PC while more than half of respondents wrongly described PC as pain medicine (60.5%) and rehabilitative medicine (57.7%). This shows that most students are unable to clearly differentiate between PC and other similar aspects of medicine which may be due to limited information resulting to inadequate knowledge. Most respondents believe that the philosophy of PC affirms life (62.6%) and recognizes dying as a normal process (69.4%). More than three-quarter of respondents believed that PC does not hasten death, although more than half wrongfully felt PC inversely prolongs life. This shows that respondents have a moderate grasp of philosophy of PC. Similarly, more than three-quarter of the respondents wrongly assumed that all dying patients need PC and felt that PC is needed for uncontrolled pain in metastatic cancer and useful in end stage heart failure while only half of the respondents felt it wasn’t needed for three days after gall bladder surgery with pain. This shows that respondents assumed PC is needed in acute postoperative pain and were unable to differentiate acute pain medicine, palliative medicine and hospice care. In the knowledge of morphine usage and its side effects, more than half of respondents believed morphine improves quality of life while 12.1% felt morphine causes death in all dying patients. Majority of students (77.2%) had a false opinion of morphine to be a universal analgesic for all kinds of pain and only 35.6% of students knew about morphine’s role in relieving breathlessness in heart failure. Students acknowledged nausea and vomiting (57.7%) and drowsiness (78.3%) as common side effects of morphine in PC setting while 56.6% were unaware of constipation and a significant number of students (78.6%) were wrong about morphine causing addiction. This showed considerable deficiency in knowledge of opioids and analgesic drugs such as morphine among clinical physiotherapy students in Nigeria. This may be due to poor background knowledge of pharmacology received by physiotherapy students at undergraduate level.

Furthermore in this study, over half of respondents could appropriately identify non pain symptoms in PC settings and majority correctly identified components of good death. A good number of physiotherapy students have better knowledge of resuscitation in PC as compared to previous study that assessed PC knowledge among other healthcare students in India by Sadhu et al., [12]. Over two-third of respondents think patients and relatives should always be involved in decision making of ―not for resuscitation‖. Almost half of the students (49.1%) correctly perceive that the statement ―Nigerian legal system at present has laid down clear guidelines about end of life and not for resuscitation‖ is false. This uncertainty in knowledge by half of respondents might be as a result of lack of exposure to the medical policies of the Nigerian legal system and the ineffectively implemented policies put in place by the Nigerian government concerning PC as reported by Chukwunyere, [9]. When communicating prognosis, a good majority of students agreed that prognosis(92.2%), patient’s wishes and choices(92%) should be clearly conveyed and that not communicating prognosis could lead to lack of trust (84%), however, 43.1% felt prognosis should only be informed to family members. This showed that students have a good understanding of communicating prognosis in PC and results might be influenced by a mild sense of empathy. Majority of students correctly identified medical social worker, nurse and occupational therapist as part of PC multidisciplinary team but also included radiotherapist who is a seldom part of multidisciplinary PC team as noted in the study by Sadhu et al., [12]. This inclusion may be as a result of misunderstanding or inexplicit knowledge in differentiating between the roles of a radiologist and a radiotherapist. Favourable attitude of almost all respondents towards PC may indicate that many students were uncertain about a number of choices and the most preferred choices still had below average ratings. This finding is fairly consistent with earlier study by Sadhu et al., [12] which suggested that students were still uncomfortable in facing matters of death and dying. This study however showed that some students displayed a certain sense of empathy and may be able to deal with PC in a professional and sensitive manner that meets the needs of both patients and family.

University of respondents was the only socio-demographic variable that had influence on his/her knowledge of PC. This may be due to factors such as different teaching methods adopted by universities, quality and quantity of teaching, available human, capital and material resources, complete absence or deficiency in PC curriculum offered by universities which births a need to review or upgrade the PC curriculum in Nigerian universities. Respondents’ attitude towards PC was also influenced by each of age and tribe of respondents. The influence of age on PC may indicate a difference in maturity level among respondents as more matured respondents may have more exposure or understanding of matters pertaining to dying and death and may cope better facing death and dying. This inference is in comparison with a study conducted at Obafemi Awolowo University on Nurses attitude towards caring for dying patients in a Nigerian teaching hospital by Faronbi et al., [21] where age was a major influencing factor of attitude. Belonging to an ethnically and culturally diverse country like Nigeria can also have a significant influence on respondents attitude towards PC as some cultures may have certain beliefs regarding death and dying which are passed down to offspring. For instance, it may be considered inappropriate, insensitive or a bad omen to discuss impending death in a tribe or culture while many cultures do not accept or encourage the use of opioids, which are the primary pharmacologic treatment for pain relief PC [22]. This shows how much of a challenging barrier, age and ethnicity or tribes can have on the attitude and invariably knowledge of respondents towards PC. Lastly, respondents’ knowledge influenced their attitude towards PC and vice versa. This may imply that having above average knowledge of PC in this sample of respondents resulted in favourable attitude towards PC.

In conclusion, there are some deficiencies and inadequacies in basic aspects of PC knowledge among clinical physiotherapy undergraduates in Nigeria. Respondents however demonstrated positive attitudes towards PC. Respondents’ knowledge and attitude were influenced by university attended which may be a factor of quality and quantity of teaching, available resources and personnel; age which may indicate maturity, individuals’ level of exposure and life experiences as well as geopolitical zone of origin, which may be a reflection of cultural beliefs and practice. These indicate an essential need for inclusion, reform or improvement in PC in the curriculum of physiotherapy undergraduate training in Nigeria.

This study has some limitations. The self-report mode of questionnaire administration used to assess the knowledge and attitude of respondents might have subjected it to response bias, so the researcher had to assume that the respondents were truthful. This study was also limited by the small sample of respondents in some of the universities with larger student population thereby skewing towards some Universities than others. The results may not indicate a true measure of knowledge and attitude of students in some Universities like Bayero University, Kano with a comparatively small number of respondents in relation to student population.

Results of this study should be interpreted with caution. Over 40% of respondents were from one institution, Obafemi Awolowo University, southwest Nigeria while samples from other Universities appeared disproportionate to eligible students’ population. Noteworthy is the comparatively small number of respondents from Bayero University, Kano northeastern Nigeria, and Nnamdi Azikiwe University, Anambra southeastern Nigeria, despite comparatively high number of eligible student population with Obafemi Awolowo University. This may imply that results of this study are dependent more on the knowledge of respondents from southwestern Nigeria.

This study was conducted among clinical physiotherapy undergraduates across Nigerian universities where Physiotherapy is offered as a course of study. Although each university has different quota based on many factors, respondents in this study received training under similar conditions and represent the Nigerian students. Hence the result of this study can be generalized to Nigerian Physiotherapy students.

## Acknowledgments

We thank all undergraduate clinical physiotherapy students in all Nigerian Universities who partook in this study. We are also grateful to the executives of National Association of Physiotherapy Students for facilitating meeting with student representatives of each University

## Author Contributions

M.O.O conceptualized the study, conducted the research, prepared the manuscript, and took part in data analysis and interpretation. J.C.N was involved in conceptualization of the study, data collection, provision of study material and preparation of manuscript. A.M.O and O.I.A were involved in the research design, administrative support and proofread the manuscript for intellectual contribution. A.M.O assembled the data, was involved in data analysis and interpretation, and proofreading. All authors read and approved the final manuscript.

### Data Citation

Data generated from this study are available on reasonable request from the corresponding author

### Ethical statement

The authors are accountable for all aspects of the work in ensuring that questions related to accuracy or integrity of any part of the work are appropriately investigated and resolved. Written informed consent was obtained before data collection. This research was approved by Health and Research Ethics Committee of Institute of Public Health, Obafemi Awolowo University, Ile Ife Nigeria with protocol number ERC/2021/05/10. Consent for publication is not applicable to this study as no identifiable details of respondent was collected or published.

**APPENDIX I:** Structured questionnaire for assessment of Palliative care knowledge as used by Kassa et al., 2014.

| Item no. | Question | Correct answer |
| --- | --- | --- |
| A) | **PC is**<br>Pain medicine | No |
|  | Geriatric medicine | No |
|  | Rehabilitation medicine | No |
|  | Active care of the dying | Yes |
| B) | **Philosophy of PC**<br>Affirms life | Yes |
|  | Recognizes dying as a normal process | Yes |
|  | Hastens death | No |
|  | Prolongs life | No |
| C) | **PC is needed for**<br>All dying patients | No |
|  | Metastatic cancer with uncontrolled pain | Yes |
|  | End stage heart failure | Yes |
|  | Three days after gall bladder surgery with pain | No |
| D) | **Morphine**<br>Causes death in all dying patients | No |
|  | Improves quality of life | Yes |
|  | Relieves all kinds of pain | No |
|  | Relieves breathlessness in heart failure | Yes |
| E) | **Common side effects of morphine in PC setting**<br>Nausea and vomiting | Yes |
|  | Constipation | Yes |
|  | Drowsiness | Yes |
|  | Addiction | No |
| F) | **Common non-pain symptoms encountered in PC**<br>Delirium | Yes |
|  | Constipation | Yes |
|  | Vomiting | Yes |
|  | Breathlessness | Yes |
| G) | **Components of good death**<br>Pain and symptom management | Yes |
|  | Clear decision making | Yes |
|  | Preparation for death | Yes |
| H) | **Resuscitation in cancer patients**<br>Resuscitation is always appropriate in advanced metastatic cancer. | No |
|  | Outcomes of resuscitation in advanced metastatic cancer are always good. | No |
|  | Patients and relatives should always be involved in decision making of “not for resuscitation”. | Yes |
|  | Nigerian legal system at present has laid down clear guidelines about end of life and not for resuscitation. | No |
| I) | **Communicating prognosis**<br>Prognosis should always be clearly communicated. | Yes |
|  | Prognosis should only be informed to family members. | No |
|  | Not communicating prognosis could lead to lack of trust. | Yes |
|  | Patient’s wishes and choices should be clearly communicated. | Yes |
| J) | **PC multidisciplinary team comprises of**<br>Medical social worker | Yes |
|  | Nurse | Yes |
|  | Radiotherapist | No |
|  | Occupational therapist | Yes |

**Appendix II:** FROMMELT attitude towards care of the dying scale (FATCOD)

| Item no. | Question |
| --- | --- |
| 1. | Giving care to the dying person is a worthwhile experience. |
| 2. | Death is not the worst thing that can happen to a person |
| 3. | I would be uncomfortable talking about impending death with the dying person. |
| 4. | Caring for the patient's family should continue throughout the period of grief and bereavement. |
| 5. | I would not want to care for a dying person. |
| 6. | The non-family caregivers should not be the one to talk about death with the dying person. |
| 7. | The length of time required to give care to a dying person would frustrate me. |
| 8. | I would be upset when the dying person I was caring for gave up hope of getting better. |
| 9. | It is difficult to form a close relationship with the dying person. |
| 10. | There are times when death is welcomed by the dying person. |
| 11. | When a patient asks, "Am I dying?" I think it is best to change the subject to something cheerful. |
| 12. | The family should be involved in the physical care (feeding, personal hygiene) of the dying person. |
| 13. | I would hope the person I'm caring for dies when I am not present |
| 14. | I am afraid to become friends with a dying person. |
| 15. | I would feel like running away when the person actually died. |
| 16. | Families need emotional support to accept the behaviour changes of the dying person. |
| 17. | As a patient nears death, the non-family caregiver should withdraw from his or her involvement with the patient. |
| 18. | Families should be concerned about helping their dying member make the best of his or her remaining life. |
| 19. | The dying person should not be allowed to make decisions about his or her physical care. |
| 20. | Families should maintain as normal an environment as possible for their dying member. |
| 21. | It is beneficial for the dying person to verbalize his or her feelings. |
| 22. | Care should extend to the family of the dying person. |
| 23. | Caregivers should permit dying persons to have flexible visiting schedules. |
| 24. | The dying person and his or her family should be the in-charge decision makers. |
| 25. | Addiction to pain relieving medication should not be a concern when dealing with a dying person. |
| 26. | I would be uncomfortable if I entered the room of a terminally ill person and found him/her crying |
| 27. | Dying persons should be given honest answers about their condition. |
| 28. | Educating families about death and dying is not a non-family caregiver's responsibility. |
| 29. | Family members who stay close to a dying person often interfere with the professional's job with the patient. |
| 30. | It is possible for non-family caregivers to help patients prepare for death. |

